# SOM-based embedding improves efficiency of high-dimensional cytometry data analysis

**DOI:** 10.1101/496869

**Authors:** Miroslav Kratochvíl, Abhishek Koladiya, Jana Balounova, Vendula Novosadova, Radislav Sedlacek, Karel Fišer, Jiří Vondrášek, Karel Drbal

## Abstract

Efficient unbiased data analysis is a major challenge for laboratories handling large flow and mass cytometry datasets. We present EmbedSOM, a non-linear embedding algorithm based on FlowSOM that improves the analysis by providing high-performance embedding method for the cytometry data. The algorithm is designed for linear scaling with number of data points, and speed suitable for interactive analysis of millions of cells without downsampling. At the same time, the visualization quality of single cell distribution within cellular populations and their transition states is competitive with the current state-of-the-art algorithms. We demonstrate EmbedSOM properties on two use-cases, showing benefits of using the interactive algorithm speed in supervised hierarchical dissection of cell populations, and the scalability improvement by efficiently processing very large datasets.

## 1. Introduction

The ever-increasing size and dimensionality of data generated by flow and mass cytometry experiments drive interest in simplifying data analysis. Employing the usual repetitive manual gating, exploration and back-gating techniques is tedious if the sample count is high, and becomes imprecise on complex data. During the past decade, a multitude of automated analysis methods have been introduced, including various unsupervised clustering and phenotyping algorithms, and embedding methods. Comprehensive reviews of the algorithms are available [19, 27, 10, 11].

The preferred method to *display* cytometry datasets is embedding, in which cells are arranged into a 2-dimensional picture showing populations of agglomerated cells with similar properties. This provides a straightforward way to inspect the relative population sizes, their contents, and the presence of various features including major subpopulations, intermediate cell states and trajectories of their development. The performance of available embedding algorithms is constantly being improved. For example, the widely used tSNE [24] has formed the basis for faster ASNE [17] and HSNE [16], and was further improved in OptSNE and FItSNE [3, 14] and accelerated using GPU by Chan et al. [5]. Other methods include SWNE, scvis, largevis [29, 6, 8], and the relatively new UMAP [15] that specifically aims to provide better, faster embedding than tSNE. Despite these developments, two key objectives have not been met:

- The embedding algorithm should be able to process the data of volumes common in cytometry quickly, ideally within seconds, to allow interactive data inspection;
- algorithm processing time should scale linearly with the number of cells, to be able to keep up with the increasing sizes of datasets without down-sampling.

We introduce EmbedSOM, a new embedding algorithm that is designed to satisfy these two requirements. The algorithm uses a self-organizing map (SOM) that describes the multidimensional cell space. SOMs have been successfully used for classifying this space into clusters – for example, FlowSOM [25] uses SOM vertices as cluster centers to classify the cells into clusters that form a basis for further analysis. The SOM additionally approximates a section of a *smooth* 2-manifold embedded in the multidimensional space in a manner such that the cells are uniformly distributed in its neighborhood. EmbedSOM uses this additional information to compute the embedding by fitting a projection of each cell onto this manifold, and transforming the projection coordinates to 2-dimensional SOM-relative coordinates.

The performance-oriented design of EmbedSOM differs substantially from other commonly used embedding methods. Most importantly, the usual time-consuming iterative optimization of single cell positions in the embedding is replaced by relatively fast manifold approximation by SOM training. Additionally, the separation of the SOM-training from other stages of the algorithm introduces flexibility that allows precise manipulation of separate embedding properties, such as reliable alignment of cell populations in the embedding even with 60 substantially different samples, and easy optimization of the embedding layout.

Here, we describe the properties of the EmbedSOM algorithm, mainly as results of benchmarking it against other embedding algorithms on several datasets. In addition to evaluating performance, our benchmark also assessed how well the embedding displays different cell populations present in manual gating of published datasets. We further demonstrate the performance properties of EmbedSOM on two analyses of large multi-sample datasets. In the first, the embedding is used to provide a visual guide for interactive FlowSOM-assisted cell population dissection, which makes the analysis more accessible for scientists, and vastly improves both precision and data throughput when compared with traditional manual gating. In the second, we demonstrate the scalability and resource efficiency by embedding and analyzing a dataset of 24 million cells. In both cases, we use the connection to the underlying FlowSOM information to provide a synoptic view of the multi-sample population differences.

## 2. Results

### 2.1. EmbedSOM provides superior embedding speed

The main distinctive feature of EmbedSOM is its computational efficiency. A dataset of common size (approximately 300k cells and 20 markers) can be mapped by the SOM and embedded in less than a minute; the GPU-accelerated versions of the algorithms deliver the same result in seconds. Generally, embedding datasets with millions of cells and several dozen markers is possible in minutes using common office hardware. Moreover, since the major SOM-training part of the required computation is shared with FlowSOM, EmbedSOM visualization adds only minor computational complexity to workflows that already use FlowSOM.

Qyantitative measurements of the speed advantage on the benchmark computations are displayed in Figure 1a. The results confirm the expected performance scaling gap between UMAP and EmbedSOM, caused by different algorithm design. Generally, UMAP and tSNE are very efficient for high-dimensional data with a low data point count, because they remove the dimensionality overhead early in the process but require more than linear amount of computation to optimize the final data point positions. The performance of both EmbedSOM stages is linear in the number of data points, but the dimensionality overhead is present throughout almost the entire computation as another linear factor.

**Figure 1:**
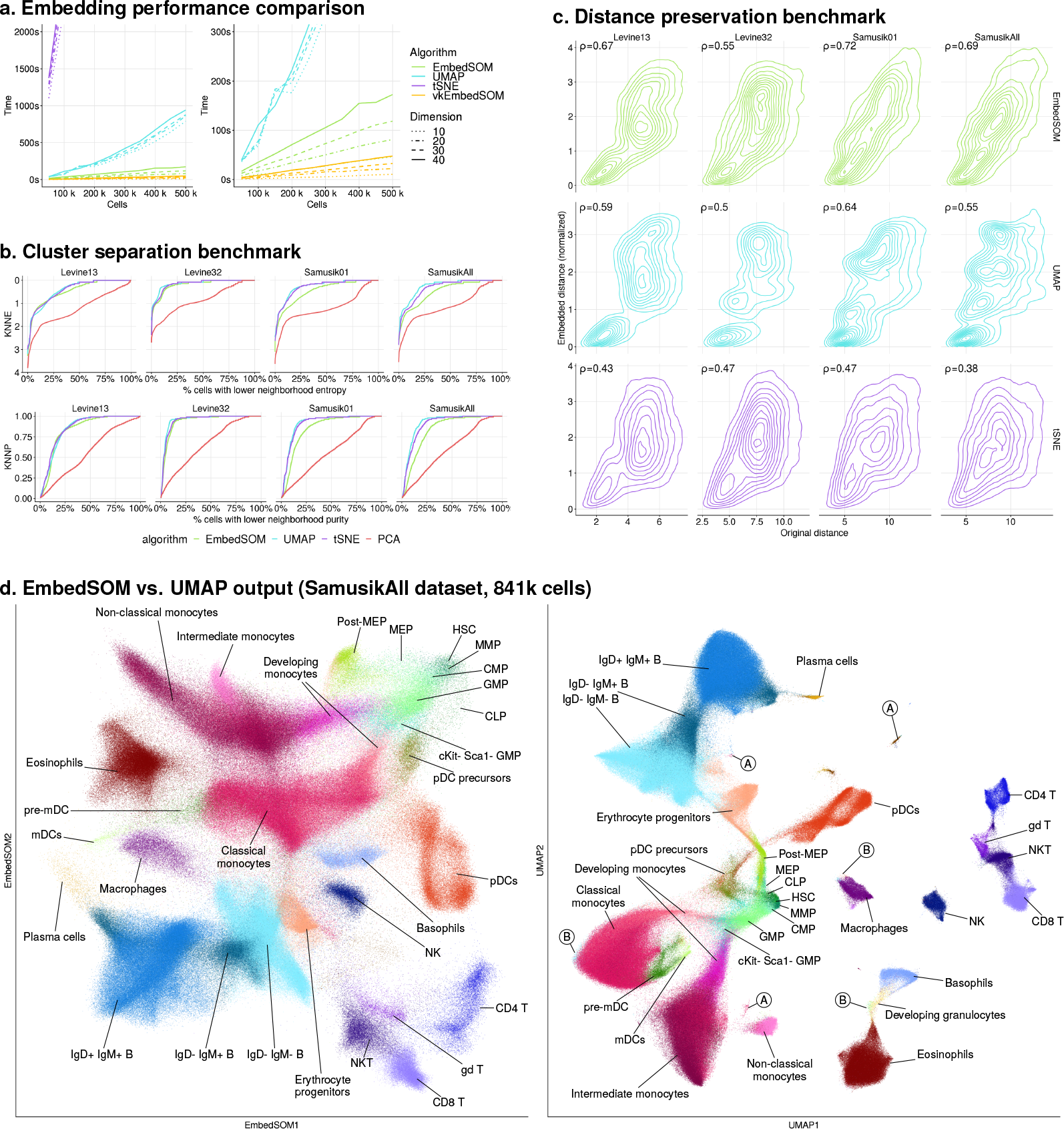
Benchmark of EmbedSOM properties compared to other embedding methods. **a.** Performance comparison, purposefully plotted using linear scales to aid realistic depiction of speed and scaling difference. **b.**,**c.** Comparison of qualitative properties of embedding. Values of Pearson correlation of original vs. embedded distances are reported in corresponding plots in **c**. **d.** Side-by-side comparison of EmbedSOM and UMAP output on SamusikAll dataset. Both methods escaped planar topology limitations by bridging clusters (developing erythrocytes over monocytes in case of EmbedSOM, and over developing pDCs in case of UMAP). High-variance cell distributions are displayed differently — EmbedSOM retains relative variance in the sample, UMAP creates population-like cluster artifacts (labeled with A) or attaches the scattered cells to a larger cluster (B). EmbedSOM calculation took 2.5 minutes, giving 15× speedup over 38 minutes required by UMAP.

EmbedSOM benefits from this trade-off when applied to high-volume flow and mass cytometry data that are physically limited to dozens of dimensions. Conversely, UMAP remains faster on low-volume datasets with more than hundreds of dimensions. For example, the raw single-cell RNA sequencing data must be pre-processed (e.g. reduced to principal components) in order to deliver comparable results with SOM-based algorithms [7].

Overall, the performance scaling difference is best illustrated by the computation time required for running workflows used later in the article: Embedding-assisted dissection of 1 million cells (Section 2.3) takes less than 5 minutes of computation with EmbedSOM; but more than 1 hour with UMAP. Embedding of the 24 million cells from Pregnancy dataset (Section 2.4) can be finished faster than in 1 hour with EmbedSOM, but requires around 2 days with UMAP.

### 2.2. EmbedSOM visualization quality is competitive with other algorithms

The quality of visualization was measured by quantifying the contents of *k*-neighborhoods in the embedding, and comparing it to the published manual gating of the same datasets. Optimally, the embedding algorithms should not mix cells from different identified populations in the same area of the resulting picture. To assess that, we measured *k*-NN (*k* nearest neighbor) entropies and *k*-NN purities of the embedded benchmark datasets (see Section 4.2.3 for definitions of the measures used, and supplementary Section S4 for description of testing datasets). Results are shown in Figure 1b. All algorithms provided visualizations of comparable quality, and we consider the slight disadvantage of EmbedSOM to be a reasonable trade-off for the performance gain.

The quality difference between EmbedSOM, UMAP and tSNE, most visible in the more interspersed part of the data, arises from the design of the algorithms. Specifically, neither UMAP nor tSNE aim to preserve local linearity of the transformation, which allows them to take apart the clusters with noisy data and attach the residual noise to nearest clusters. This makes the embedding arguably more visually appealing by creating well-defined, undistorted borders, and at the same time improves the *k*-NN-based measures by reducing the chance of a cell from a different population occurring in a *k*-neighborhood.

Figure 1c shows comparison of original vs. embedded distances of cell pairs, which can be used to interpret how faithfully the algorithms reproduce small- and large-scale distances in the data. This measure has also been used by Becht et al. [2]. Results are summarized as Pearson correlations of original and embedded distances, labeled as *ρ* in each plot.

The distance preservation plots show one possible origin of the worse performance of EmbedSOM in *k*-NN-style metrics, most visible on Samusik datasets. Optimization of single cell positions used by tSNE and UMAP necessarily produces better separation of the complicated cluster structure, at the cost of introducing high quantization (in case of UMAP) and scattering (tSNE) of the embedded distances.

While the observed cluster separation may be desirable if the embedding is expected to approximate the population boundaries, it may be inappropriate if the population environment is relevant for analysis. For example, tight packing of cells impairs the possibility to observe the natural population density distribution or to filter out noise manually. The differences can be clearly observed in the embeddings of the complete SamusikAll dataset in Figure 1d. The present artifact clusters (marked with A) and cells attached to borders of other clusters (marked with B) are highly undesirable in many situations, such as analysis of development trajectories.

On the other hand, compaction of the noise can bring up various high-variance data, such as the population of developing granulocytes displayed by UMAP in Figure 1d. EmbedSOM displays this population correctly as connecting the progenitors with basophils and eosinophils without the gap (neutrophils are not present since they are excluded from SamusikAll dataset), but the limitations of planar topology cause it to be displayed in overlay with other populations and thus not easily visible (see Figure S4).

Such properties of the algorithms are best interpreted as different balances in failure modes, which can guide the choice of optimal algorithm for specific visualization requirements: Compacting the residual or unexplained noise is desirable for providing a clean display of the data for publication. On the contrary, almost-immediate availability of all information about very large datasets, including the (often informative) noise, is more important for producing comprehensive graphics for high-throughput analyses.

### 2.3. Fast embedding augments supervised hierarchical dissection analysis

Manual and supervised analysis tools and algorithms are usually required to provide quick feedback to user actions, in order to be practical and avoid unnecessary workflow delays. Delivering the embedding performance sufficient for use in interactive tools is the main aim of EmbedSOM. In this section, we show that EmbedSOM integrates into typical supervised analysis and improves its throughput and precision.

For demonstration, we augment the usual hierarchical gating-like dissection by using results from high-dimensional unsupervised analysis. While embedding has been already used for hierarchical sample dissection [23] as a way to provide highly informative and comprehensible overview of the high-dimensional data that can precisely guide the analysis, EmbedSOM improves the analysis performance by removing unnecessary delays and interruptions required for computation on large datasets. Moreover, quickly interleaved user input and computation phases facilitate easier and faster exploration of the datasets.

The workflow is constructed as follows: FlowSOM is used internally for clustering the cells, but the user is only presented with the results in EmbedSOM embeddings, which provides a more cell-centric way to look at FlowSOM output. The presented layout of cell populations corresponds to the positions of the clusters in the grid-view output of FlowSOM [25, Fig. 1 (ii) and 2], but, as a main improvement, the cells are distributed in a way that retains the local topology and variance in the sample. Consequently, the population shapes and density distributions (which would otherwise disappear in the view of FlowSOM star plots) are observable similarly as in the usual two-dimensional dot-plots.

Cell population dissection is realized by user selection of FlowSOM metaclusters from the embedding. That is both more natural to scientists who are used to the prevalent dot-plot view, and less prone to various statistical and bias-derived errors, since human choice is restricted to a *discrete and reproducible* selection of FlowSOM clusters that have previously been proven to capture the respective cell populations very precisely [27]. The selected metaclusters are used either as final identified populations, or as a basis for further hierarchical dissection process that continues by re-clustering and re-embedding of the selected cell subset (see Section 4.2.4).

We used this workflow to dissect samples from a study of newly generated transgenic mouse model that was screened for changes against the respective wild-type. The workflow is displayed in Figure 2: First, all compensated samples (*n* = 11, see Section S1 for animal details and staining) from a 14-dimensional dataset were aggregated and subjected to clustering and embedding. The user was let to choose clusters of interest from the view of all events (Figure 2a), which was used to remove the dead cells and debris to get a clean picture of a subset of live cells (Figure 2b), then to further refine the selection to T cell and NK cell populations (Figure 2c), and, following the gating hierarchy from Figure S1, to display the CD4 T cell population (Figure 2d). The last level uses the MST-based embedding layout for improved cluster separation (see Section S2).

**Figure 2:**
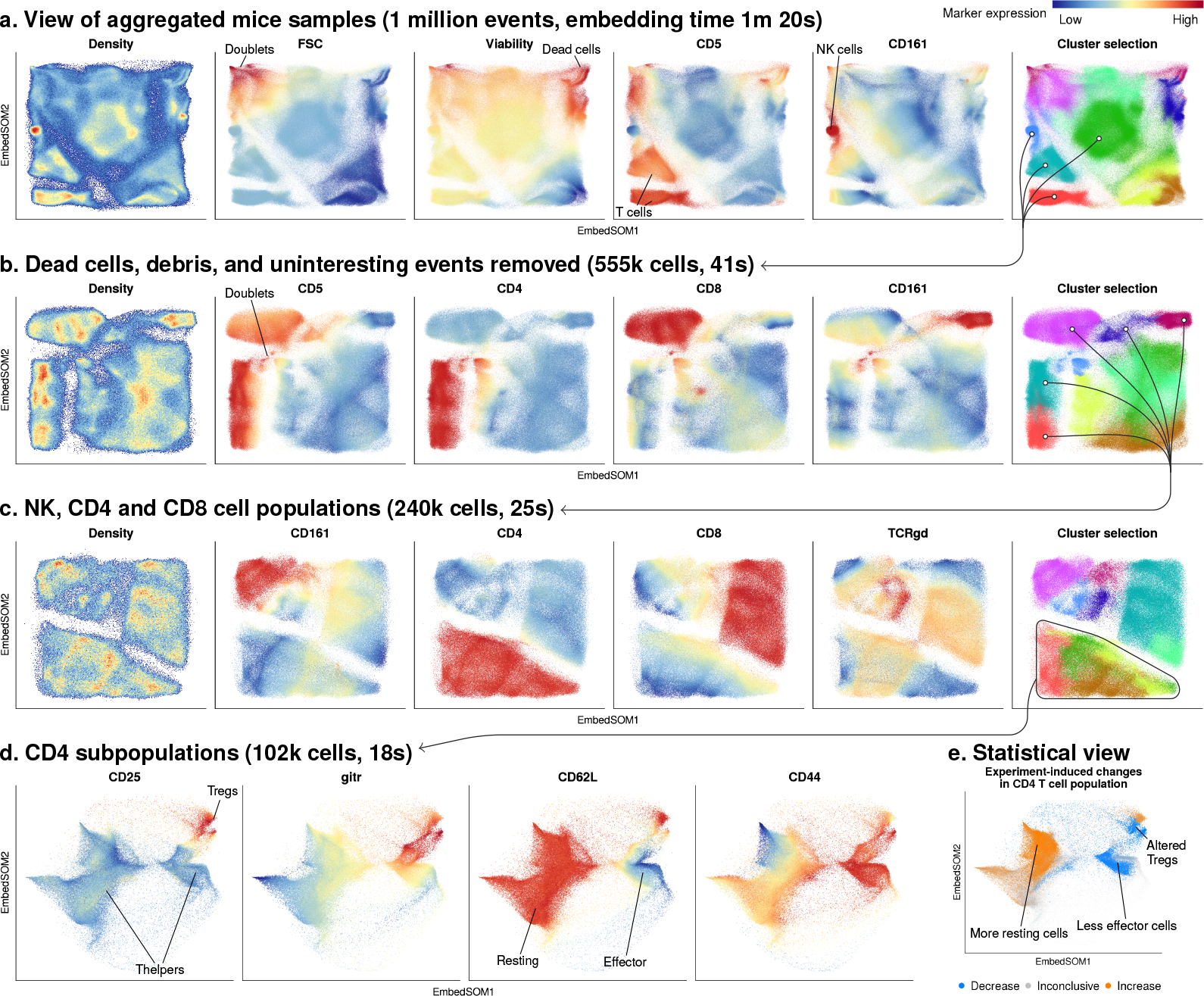
Hierarchical sample dissection assisted by EmbedSOM view, as described in Section 2.3. Total run times for EmbedSOM and FlowSOM metaclustering are reported in parentheses. The user advanced from the view of all cells (**a**) by progressively reducing the examined cell subset to a selection of metaclusters generated at each level (the selection is displayed as arrows). This was used to remove dead cell and debris (**b**) and reduce the live cells to T and NK cells (**c**). Internal distribution of CD4 T cell subpopulations (already noticeable in the density plot in **c**) is further explored in the last levels of dissection (**d**). Direct visualization of statistical testing results of the demonstration experiment (**e**) shows color-coded changes of cell count in respective subpopulations.

After the dissection, the identified subpopulations were used for statistical testing of the experiment outcome. For that purpose, we exploited the correspondence of EmbedSOM embedding with the underlying FlowSOM cluster structure to display FlowSOM-derived statistical information directly in the embedding. In particular, we found it practical to quickly display the significant changes in the contents of FlowSOM clusters by color-coding the embedding with the results of bulk statistical testing, as seen in Figure 2e. This information can be used to guide the precise statistical testing accordingly (see Section 4.2.4 for description).

On the testing laptop, the total computational time required to process the aggregated sample of 1 million cells was less than 4 minutes, which we consider to be a reasonable overhead for interactive workflow. The whole EmbedSOM-assisted analysis, including the time required for human input, was finished in a fraction of time that was required for the corresponding manual dissection, and resulted in the same observation of decrease in effector CD4 T helper cells (*p* ≤ 0.05 in manual analysis), increase in resting CD4 T helper cells (*p* ≤ 0.08), decrease in effector C4 Treg cells (*p* ≤ 0.02) and increase in resting CD4 Treg cells (*p* ≤ 0.02).

### 2.4. High-throughput embedding allows improved statistical display of data

The original aim of EmbedSOM was to simplify visualization of differences in high-volume multi-sample data.

We demonstrate this functionality in Figure 3, on a mass-cytometry dataset from recent study by Aghaeepour et al. [1]. The data were collected from 18 women where the whole-blood samples were collected in 4 time points during and after pregnancy (early, mid- and late-pregnancy, and 6 weeks postpartum). All samples were measured in unstimulated state as well as stimulated with LPS, IFN-*α* and IL-2+IL-6, with total 36 markers used for studying immune system function and regulation. A selection of 112 samples used for this demonstration (cca. 39 million cells) was gated to single/leukocyte/non-erythrocyte/non-platelet; the final dataset contained approximately 24 million cells.

SOM-style analysis and embedding can process this amount of cells efficiently without any required downsampling. The nature of both EmbedSOM stages allow the data to be processed as a stream; consequently, memory requirements of SOM training and embedding are constantly low, regardless of dataset size. Exploiting this data streaming technique, we were able to embed all 24 million cells in under 1 hour on an office-grade laptop computer.

**Figure 3:**
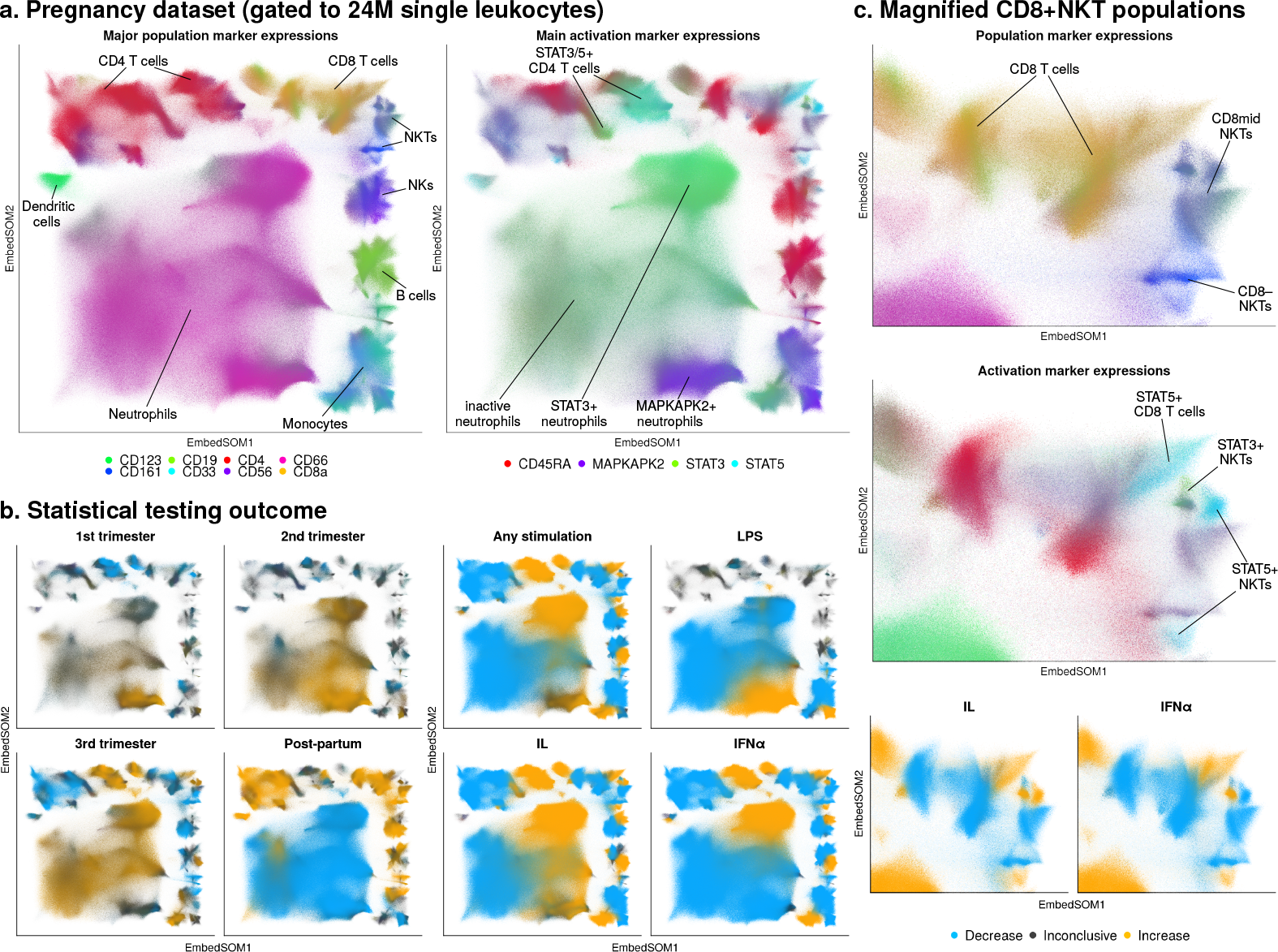
Application of EmbedSOM to large data. **a.** Selection of surface marker expression and signaling molecule activation displayed the embedding of 24 million cell events in Pregnancy dataset. Colors are blended as gradients; cells that do not express any selected markers are displayed as gray. **b.** High number of processed cells allows more precise statistical testing on all subpopulations: dataset is displayed at different time points (left) and different stimulations at 3^rd^ trimester (right) compared to all other respective states and colored accordingly to the significance of the difference. **c.** Magnified view of the upper-right corner of the whole embedding reveals small changes in the CD8 and NKT populations.

Technically, all cells were processed individually, but the design of the algorithm guaranteed correct positioning of the cells in the embedded populations. This property of EmbedSOM can be used to reliably produce correctly aligned embeddings even from highly different cell samples.

High cell count in the whole dataset is beneficial for obtaining precise results from statistical hypothesis testing. We have applied this testing on the contents of FlowSOM clusters, and highlighted the statistically significant changes in the embedding. Similar bulk visualization of the cluster differences has been already used in other algorithms, such as diffcyt [26, Figure 2c,g] and Cytofast [4, Figure 2]. Our approach additionally exploits the aforementioned correspondence between FlowSOM mapping and EmbedSOM embedding, which improves the presentation of results by directly connecting the statistical information with respective small areas in the embedding. In result, the plotting method produces a quickly comprehensible output regardless of the of the high number of individual statistical tests.

The resulting significance plots of the Pregnancy dataset (Figure 3b) allow straightforward observation of relative sample changes, and the plots of time development and stimulation-induced changes reproduce various findings from the original article [1]: The plot of time development shows the rapid post-partum decline of subpopulations of neutrophils, additionally it identifies a similar trend in monocytes. Stimulation-induced changes are clearly observable in cell populations of CD4 and CD8 T cells, which move to corresponding STAT3^+^or STAT5^+^ regions (in agreement with the results of Tanabe et al. [20]), and in populations of neutrophils and monocytes, which, depending on the type of stimulation, move accordingly to MAPKAPK2^+^ or STAT3^+^ regions.

Individual cells start to become apparent only after magnifying the embedding in Figure 3c, which also gives a clear view of small NKT cell activation differences in IL-2+IL-6 and IFN-*α* stimulation.

## 3. Discussion

EmbedSOM alleviates the long-standing unavailability of scalable, fast non-linear embedding algorithm for single-cell cytometry data. In its current version, it extends the usefulness of commonly available hardware for running analyses and producing visualizations of high-volume datasets. Here, we draw attention to algorithm and benchmark details, and possible directions for future research.

### 3.1. Benchmarking methodology

Currently, there is no single accepted methodology for benchmarking the quality of embedding algorithms. In this work, we use the benchmark measure of *k*-NN population label entropies and purities (see Section 4.2.3 for definition). Other available measures include e.g. the Kullback-Leiber (KL) divergence of distance distribution, various measures derived from similarity between *k*-neighborhoods of a data point in high-dimensional vs. embedding space (used e.g. by Becht et al. [2]), and the NPE and residual variance (used by Konstorum et al. [11]).

We made the choice of the *k*-NN-based purity and entropy measures due to our focus on high-throughput cytometry data, for which neither the KL divergence minimization nor *k*-neighborhood preservation is the primary desired feature. Cells typically form relatively dense populations of a single cell type with broad range of accepted difference in specific gene expression (e.g., cells with a 0.1% difference in marker expression are usually considered the same). Therefore, algorithms should aim to separate the cells that are expected to belong to different populations, which is described by the *k*-NN measures, rather than attempt to preserve the irrelevant inner structure of such populations, which is required to produce a high similarity of *k*-neighborhoods and low KL divergence.

### 3.2. Hierarchical dissection of cell populations

The workflow demonstrated in Section 2.3 serves as a valuable alternative to the commonly used manual gating strategies. Apart from the improvement in precision and reproducibility, the computational assistance avoids the need to manually draw gates, which is especially convenient if applied to multiple samples at once and connected with automated analysis of their properties. For instance, the significance plots (Figure 2e) provide a compound view of data from many samples aggregated in an easy-to-inspect image of relevant statistical information, which can quickly guide the statistical analysis to the most diverging parts of the data. A similar presentation of the sample statistics has already been proposed in diffcyt [26] and successfully used for prediction of responses to immunotherapy [12].

### 3.3. Trajectories and noise

The smoothness and local linearity of the EmbedSOM projection are valuable aids in visualizing transitions between different cell populations and their states. As discussed in Section 2.2, this comes as a trade-off — the embedding is unable to completely separate cell populations from surrounding noise and debris if these are not separable in high-dimensional space, but the same property causes it to never disrupt a trajectory that connects populations. This is prominent especially with MST-based EmbedSOM layouts (see Figure S3). At the same time, local linearity improves the depiction of cell density, which simplifies distinguishing populations and trajectories of interest from noise.

Computational trajectory inference is a topic of recent research [18]; in the future we aim to exploit the results of Wolf et al. [28] to improve the trajectory depiction in EmbedSOM.

Finally, even though the perception of 2-dimensional depiction and separation of individual populations is highly subjective, we believe that the resulting similarity of EmbedSOM embeddings to the usual dot-plot projections used for manual gating will simplify interpretation of the results by scientists.

## 4. Experimental procedures and methods

### 4.1. Data and software availability

EmbedSOM is available from https://bioinfo.uochb.cas.cz/embedsom. Repositories with source code are hosted by GitHub.

The benchmarking data were selected from public-domain datasets that previously have been used for benchmarking other algorithms [13]. The dataset used for demonstration of large-scale embedding and sample difference visualization (Pregnancy dataset) was taken from the study of Aghaeepour et al. [1], selected based on the availability of time-series, multi-sample data with different measured perturbations. A summary of all datasets used in this work is provided in Section S3.

For the analysis of embedding quality, we reused the manual gating provided in the Levine and Samusik datasets. The datasets were pre-processed as in the original article [13], ungated cells were excluded from benchmarking. The visual differences between resulting embedded samples can be observed in Figure S6.

The benchmark of performance scaling (Section 2.1) was run on cells and markers that were sampled randomly from the Pregnancy dataset.

Figure Figure 1d is produced from all cells (including the ungated cells) present in SamusikAll dataset. The displayed population annotations are different from the original published gating; we have extended the annotation to also provide information about cells not classified in the original analysis.

For demonstration of the hierarchical dissection technique in Section 2.3, we used original data from transgenic mouse spleens. A detailed description of the experiment and methods is available in Section S1.

### 4.2. Method details

#### 4.2.1. Embedding algorithm

The geometric interpretation of EmbedSOM is similar to elastic maps [9] or simplified topographic manifold projections [21]. The basic idea (Figure 4) is shared with ViSOM algorithm [30] — the first algorithm stage is conducted by any applicable SOM implementation (e.g. FlowSOM) to provide a map of the multi-dimensional space; EmbedSOM improves the second stage by using a smooth approximation of the 2-dimensional projection reconstructed from simpler, more robust projections to 1-dimensional affine spaces. That approach is less prone to graphical artifacts that arise from using small SOMs on topologically complicated high-dimensional data, and escapes the dimensionality-induced crowding effect by avoiding interpretation of distances in the multidimensional space.

**Figure 4:**
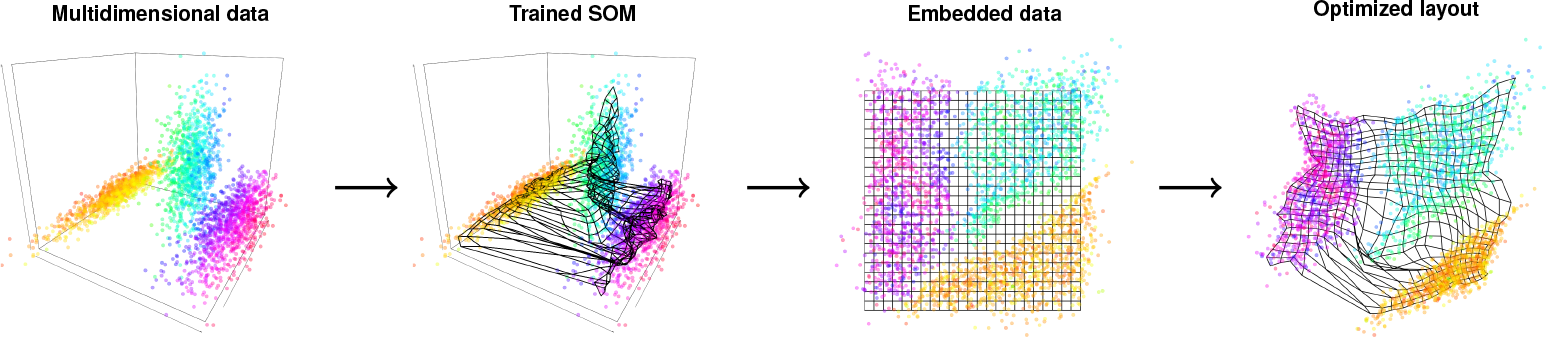
Overview of the EmbedSOM embedding process, shown on three synthetic Gaussian clusters in 3-dimensional space. From left: The cells in multi-dimensional space are used to train a SOM that describes their distribution; embedding then smoothly fits the content of multi-dimensional space to a flat version of SOM. Finally, result can be quickly optimized by changing embedding parameters and SOM layout (in the picture, the lengths of grid edges are optimized to match values in the corresponding U-matrix), and efficiently recomputing only the last algorithm stage.

The projection process in the second stage of the algorithm is separate for each cell, which directly results in the stream-processing behavior of the algorithm, and, in turn, linear computational time requirements and low memory consumption. This independence of individual cell positions in the second algorithm stage allows very simple multi-sample population alignment in the embedding; at the same time it is a basis for the efficient processing required to process datasets even larger than the 24 million cells in Section 2.4.

Mathematical description of the algorithm is available in supplementary Section S2 together with implementation details.

#### 4.2.2. Algorithm parameters

The major parameters of EmbedSOM include the SOM-training settings (shared with FlowSOM), EmbedSOM projection parameters SMOOTH, k and ADJUST, and the embedded SOM layout (see Section S2 for additional details about parameter interpretation and embedding layouts).

Correct setup of the SOM training has been previously discussed by Van Gassen et al. [25], who recommend training 10×10 SOM. We recommend using a slightly larger SOM size to provide a smoother manifold for the projection approximation. Accordingly, we used 24×24 as a default throughout this work. The SOM size should be increased for samples with large number of individual small cell subpopulations — good embedding of a population requires it to be mapped by at least one SOM vertex.

In our experience, projection parameter values of SMOOTH ∈ [−2, 2], k ∈ [15, 50] and ADJUST ∈ [0.5, 2] provided good results, and setting SMOOTH = 0, 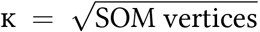, ADJUST = 1 was a good default for all data we tested. Visual differences between various parameter settings are shown in Figure S3.

For embedding, the algorithm uses a straight lattice-style SOM layout by default, but, as shown in Figure 4, any supplied layout is acceptable (examples can be found in Figures S3 and S6). In fact, Figure 1d uses this possibility to optimize the SOM layout to better match the distances in U-matrix of the SOM [22] and improve the visual quality of the result; Figure 2d uses a layout similar to the MST display as known from FlowSOM [25, Figure 4], to provide well-separated display of compacted CD4 subpopulations. Because of the small amount of data involved in the layout optimization (usually at most hundreds of SOM vertices), even very sophisticated layout algorithms may be used for the purpose with negligible impact on embedding performance.

#### 4.2.3. Benchmark setup

In the benchmark, the manual gating information required for obtaining the KNNE and KNNP metrics was obtained from publicly available expert manual gating. We measured the entropy and purity of the cell population labels (i.e. manually assigned populations) in the *k*-nearest neighborhoods of the embedded cells.

First, the data from each dataset were embedded by tSNE, UMAP and EmbedSOM, using information from all relevant markers. To compare with a linearity-preserving method, we also calculated 2-dimensional PCA projections.

We defined *k*-NN entropy as the standard information entropy of the population label values in a *k*-neighborhood, and *k*-NN purity as a probability that a random cell selected from a *k*-neighborhood belongs to the same population as the neighborhood center. These measures implicitly capture the amount of high-entropy noise and number of the misplaced cells in the embedding. Both measures were calculated for all neighborhoods of size *k* = 100 in a sample of 10,000 cells from each embedding. Individual values for the cell neighborhoods were plotted as reliability distributions to aid comparison.

Comparison of original vs. embedded cell distance distributions was performed on the same sample; the methodology is the same as in other studies [2].

We used the UMAP implementation from Python package `umap-learn` version 0.3.7 and tSNE implementation from R package `Rtsne` version 0.13 with default parameters: UMAP was run with `n_neigbors=5`, `min_dist=0.1`, `n_components=2` with 200 epochs and Euclidean metric. t-SNE was run with perplexity set to 30, *θ* = 0.5, *η* = 200, on 1000 iterations with momentum scaling from 0.5 to 0.8. For EmbedSOM, we used 24×24 SOMs built with PCA-based initialization from FlowSOM (`initf=Initialize_PCA`), layout of embedding was optimized according to U-matrix (`emcoords=’som’`); other EmbedSOM parameters were left at default values.

Performance measurements were collected on an Intel^®^ Core™ i7-4790K CPU@4.00GHz and nVidia^®^ GeForce^®^ GTX 1060 GPU.

#### 4.2.4. Workflow details

Cell population dissection was performed using DiffSOM package (see Section S3) as such: At each level, the workflow-supporting software has created a 24×24 SOM on a sample from all cells analyzed at that level, embedded it to plot marker expressions, and computed FlowSOM metaclustering for user-based selection. The user was allowed to change the parameters of SOM training and embedding to optimize the view. After examining the output, the user has selected metacluster numbers for further exploration. The same analysis was then iteratively repeated for the cells in the selected subset.

For comparison, a manual data analysis of the dataset used in Section 2.3 was performed using FlowJo software (FlowJo, LLC). The full gating strategy can be viewed in Figure S1.

To generate significance plots (as seen in Figure 3b and Figure 2e), cell counts in SOM clusters in all samples were first normalized as percentages of the entire displayed population. The percentages were grouped according to the experiment (e.g. wild-type vs. transgenic samples), and the groups were subjected to two one-sided Mann-Whitney tests. The testing measured the viability of the hypotheses of lower and higher relative cell abundance in the sample groups.

The resulting pairs of *p*-values were used for color-labeling contents of the corresponding clusters in the plot.

## Acknowledgements

This work was supported by SVV project 260451. M.K. and J.V. were supported by ELIXIR CZ LM2015047 (MEYS). A.K. and K.D. were supported by the Grant Agency of the Charles University, GAUK 1610218. J.B., and R.S. were supported by RVO68378050 (CAS), LM2015040 (MEYS), OP RDE CZ.02.1.01/0.0/0.0/16 013/0001789 (MEYS and ESIF) and OP RDI CZ.1.05/1.1.00/02.0109 (MEYS and ERDF). K.F. was supported by NV18-08-00385 (AZV).

Computational resources were provided by the ELIXIR-CZ project (LM2015047), part of the international ELIXIR infrastructure.

We are extremely grateful to Vladimír Vondruš for providing invaluable insight into Vulkan^®^ API, to Alena Keprovaá for providing datasets for testing, and to Yvan Saeys and Sofie Van Gassen for benchmarking advice.

## References

[1] Aghaeepour, N., Ganio, E.A., Mcilwain, D., Tsai, A.S., Tingle, M., Van Gassen, S., Gaudilliere, D.K., Baca, Q., McNeil, L., Okada, R., et al., 2017. An immune clock of human pregnancy. Science immunology 2, eaan2946.

[2] Becht, E., McInnes, L., Healy, J., Dutertre, C.A., Kwok, I.W., Ng, L.G., Ginhoux, F., Newell, E.W., 2019. Dimensionality reduction for visualizing single-cell data using umap. Nature Biotechnology 37, 38.

[3] Belkina, A.C., Ciccolella, C.O., Anno, R., Spidlen, J., Halpert, R., Snyder-Cappione, J., 2018. Automated optimal parameters for t-distributed stochastic neighbor embedding improve visualization and allow analysis of large datasets. bioRxiv 451690.

[4] Beyrend, G., Stam, K., Höllt, T., Ossendorp, F., Arens, R., 2018. Cytofast: A workflow for visual and quantitative analysis of flow and mass cytometry data to discover immune signatures and correlations. Computational and Structural Biotechnology Journal 16, 435–442.

[5] Chan, D.M., Rao, R., Huang, F., Canny, J.F., 2018. t-SNE-CUDA: GPU-accelerated t-SNE and its applications to modern data. arXiv preprint arXiv:1807.11824.

[6] Ding, J., Condon, A., Shah, S.P., 2018. Interpretable dimensionality reduction of single cell transcriptome data with deep generative models. Nature communications 9, 2002.

[7] Duò, A., Robinson, M.D., Soneson, C., 2018. A systematic performance evaluation of clustering methods for single-cell RNA-seq data. F1000Research 7, 1141.

[8] Dzwinel, W., Wcislo, R., Matwin, S., 2019. 2-d embedding of large and high-dimensional data with minimal memory and computational time requirements. arXiv preprint arXiv:1902.01108.

[9] Gorban, A., Pitenko, A., Zinov’ev, A., Wunsch, D., 2001. Vizualization of any data using elastic map method. Smart Engineering System Design 11, 363–368.

[10] Kimball, A.K., Oko, L.M., Bullock, B.L., Nemenoff, R.A., van Dyk, L.F., Clambey, E.T., 2018. A beginner’s guide to analyzing and visualizing mass cytometry data. The Journal of Immunology 200, 3–22.

[11] Konstorum, A., Vidal, E., Jekel, N., Laubenbacher, R., 2018. Comparative analysis of linear and nonlinear dimension reduction techniques on mass cytometry data. bioRxiv 273862.

[12] Krieg, C., Nowicka, M., Guglietta, S., Schindler, S., Hartmann, F.J., Weber, L.M., Dummer, R., Robinson, M.D., Levesque, M.P., Becher, B., 2018. High-dimensional single-cell analysis predicts response to anti-PD-1 immunotherapy. Nature Medicine 24, 144.

[13] Levine, J.H., Simonds, E.F., Bendall, S.C., Davis, K.L., ad D. Amir, E., Tadmor, M.D., Litvin, O., Fienberg, H.G., Jager, A., Zunder, E.R., Finck, R., Gedman, A.L., Radtke, I., Downing, J.R., Pe’er, D., Nolan, G.P., 2015. Data-driven phenotypic dissection of AML reveals progenitor-like cells that correlate with prognosis. Cell 162, 184–197.

[14] Linderman, G.C., Rachh, M., Hoskins, J.G., Steinerberger, S., Kluger, Y., 2017. Efficient algorithms for t-distributed stochastic neighborhood embedding. arXiv preprint arXiv:1712.09005.

[15] McInnes, L., Healy, J., Melville, J., 2018. UMAP: Uniform manifold approximation and projection for dimension reduction. arXiv preprint arXiv:1802.03426.

[16] Pezzotti, N., Höllt, T., Lelieveldt, B., Eisemann, E., Vilanova, A., 2016. Hierarchical stochastic neighbor embedding, in: Computer Graphics Forum, Wiley Online Library. pp. 21–30.

[17] Pezzotti, N., Lelieveldt, B.P., van der Maaten, L., Höllt, T., Eisemann, E., Vilanova, A., 2017. Approximated and user steerable tSNE for progressive visual analytics. IEEE transactions on visualization and computer graphics 23, 1739–1752.

[18] Saelens, W., Cannoodt, R., Todorov, H., Saeys, Y., 2019. A comparison of single-cell trajectory inference methods: towards more accurate and robust tools. Nature Biotechnology April 2019.

[19] Saeys, Y., Van Gassen, S., Lambrecht, B.N., 2016. Computational flow cytometry: helping to make sense of high-dimensional immunology data. Nature Reviews Immunology 16, 449.

[20] Tanabe, Y., Nishibori, T., Su, L., Arduini, R.M., Baker, D.P., David, M., 2005. Cutting edge: role of stat1, stat3, and stat5 in ifn-*αβ* responses in t lymphocytes. The Journal of Immunology 174, 609–613.

[21] Tino, P., Nabney, I., 2002. Hierarchical GTM: Constructing localized nonlinear projection manifolds in a principled way. IEEE Transactions on Pattern Analysis and Machine Intelligence 24, 639–656.

[22] Ultsch, A., 2003. U*-matrix: a tool to visualize clusters in high dimensional data. Technical report, published by Fachbereich Mathematik und Informatik Marburg.

[23] Unen, V., Höllt, T., Pezzotti, N., Li, N., Reinders, M.J., Eisemann, E., Koning, F., Vilanova, A., Lelieveldt, B.P., 2017. Visual analysis of mass cytometry data by hierarchical stochastic neighbour embedding reveals rare cell types. Nature Communications 8, 1740.

[24] Van Der Maaten, L., 2014. Accelerating t-SNE using tree-based algorithms. The Journal of Machine Learning Research 15, 3221–3245.

[25] Van Gassen, S., Callebaut, B., Van Helden, M.J., Lambrecht, B.N., Demeester, P., Dhaene, T., Saeys, Y., 2015. FlowSOM: Using self-organizing maps for visualization and interpretation of cytometry data. Cytometry Part A 87, 636–645.

[26] Weber, L.M., Nowicka, M., Soneson, C., Robinson, M.D., 2018. diffcyt: Differential discovery in high-dimensional cytometry via high-resolution clustering. BioRxiv, 349738.

[27] Weber, L.M., Robinson, M.D., 2016. Comparison of clustering methods for high-dimensional single-cell flow and mass cytometry data. Cytometry Part A 89, 1084–1096.

[28] Wolf, F.A., Hamey, F., Plass, M., Solana, J., Dahlin, J.S., Gottgens, B., Rajewsky, N., Simon, L., Theis, F.J., 2018. Graph abstraction reconciles clustering with trajectory inference through a topology preserving map of single cells. bioRxiv 208819.

[29] Wu, Y., Tamayo, P., Zhang, K., 2018. Visualizing and interpreting single-cell gene expression datasets with similarity weighted nonnegative embedding. Cell systems 7, 656–666.

[30] Yin, H., 2008. On multidimensional scaling and the embedding of self-organising maps. Neural Networks 21, 160–169.

