## Supplementary text, tables and figures for "SOM-based embedding improves efficiency of high-dimensional cytometry data analysis"

### S1. Transgenic mice data and analysis

*Animals.* Mice were housed at the animal facility of the Czech Centre for Phenogenomics (CCP). Cyp39a1em1(IMPC)Img/Ph (referred to as Cyp39a1$^{-/-}$) mice were generated at the Targeting Unit of CCP by CRISPR/Cas9 genome editing strategy as a part of International mouse phenotyping consortium (IMPC) effort (see `https://www.mousephenotype.org/imits/mi_attempts/18619#distribution_centres`).

All experiments were approved by the ethical committee of the Institute of Molecular Genetics of the ASCR (IMG).

*Mouse splenocyte flow cytometry analysis.* Single-cell suspensions of mouse spleens were prepared using a Mouse Spleen Dissociation Kit (Miltenyi Biotec) and gentleMACS™ Dissociator (Miltenyi Biotec) according to manufacturer's instructions. After red blood cell lysis, cells were passed through a $100\mu$m cell strainer and counted. Cells were stained with Ghost dye UV450 in HBSS (Tonbo biosciences) to exclude dead cells, next, Fc receptors were blocked (2.4G2; BD Biosciences) and cells ($2 \times 10^6$ per sample) were stained with following anti-mouse antibody cocktail:

- CD5-BV421 (53-7.3),
- CD49d-BV510 (R1-2),
- $\gamma\delta$TCR-BV605 (GL3),
- GITR-BV711 (DTA-1),
- CD4-FITC (RM4-5),
- Klrg1-PerCP-Cy5.5 (2F1/KLRG1),
- CD44-PE (IM7),
- CD8a-PE-CF594 (53-6.7),
- CD161-APC (PK136),
- CD25-PE-Cy7 (PC61),
- CD62L-APC-Cy7 (MEL-14),

(all BD Biosciences, except for Klrg1, which was purchased from Biolegend).

Fluorescence data were acquired using BD LSRFortessa™cell analyzer (BD Biosciences). Total 6 control and 7 transgenic mice were measured, from that 1 control and 1 transgenic mice samples were removed during quality control as outliers.

Manual data analysis (see below) was performed using FlowJo software (FlowJo, LLC). EmbedSOM-assisted data analysis was performed using the DiffSOM wrapper[1].

*Results.* The manual gating strategy used for retrieving the cell populations in FlowJo is shown in Figure S1.

[1] `https://github.com/exaexa/DiffSOM`

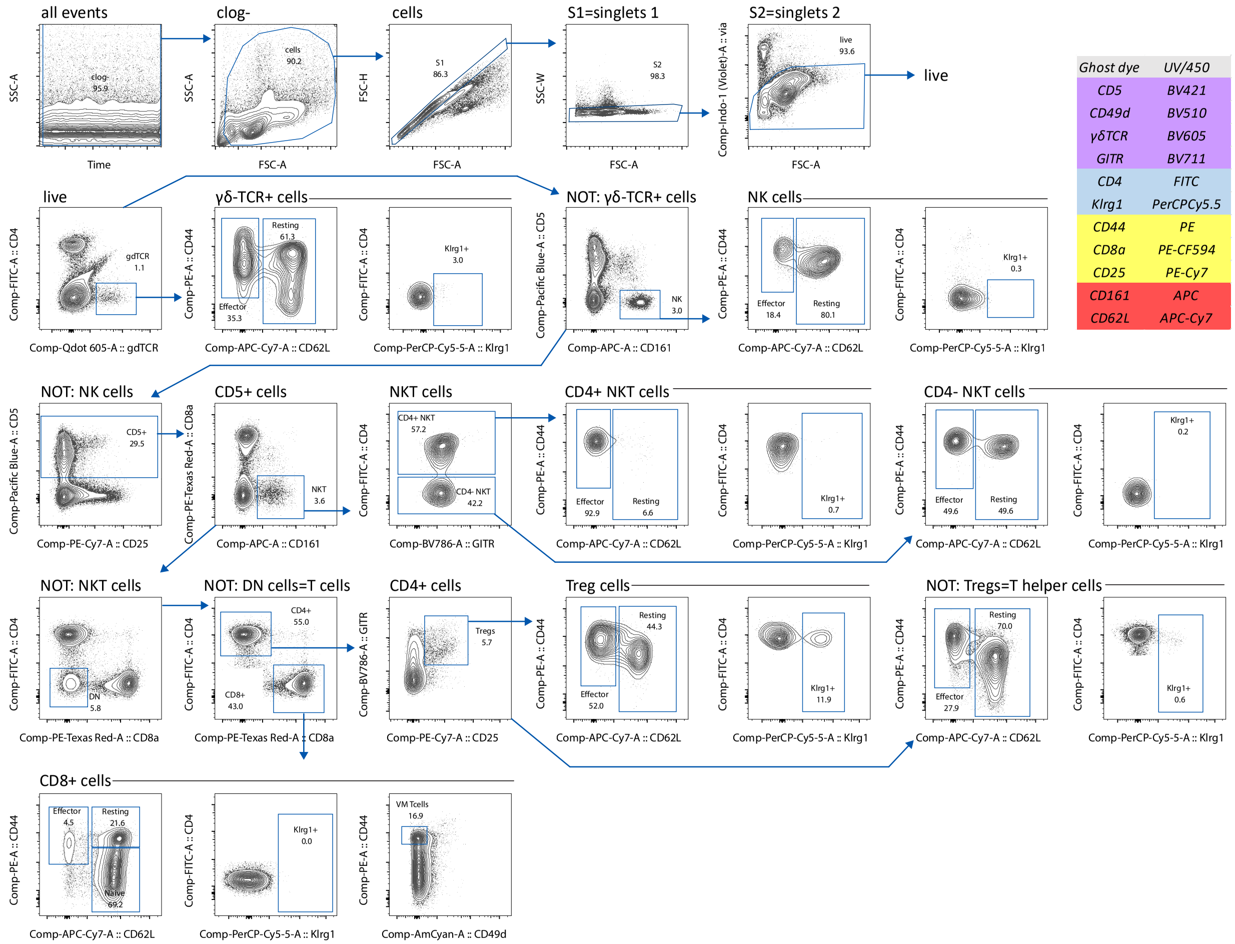


Figure S1: Gating strategy used in the manual analysis that corresponds to the analysis in Section 2.4 and Figure 2.

### S2. EmbedSOM algorithm and parameters

EmbedSOM projection process in the second stage of the algorithm is separate for each cell, and depends on the set $G$ of the positions of SOM vertices, set $G'$ of the corresponding positions of the vertices in the low-dimensional embedding space, and several tunable parameters (described below). The following description corresponds to the illustration in Figure S5.

To embed the cell at position $c$, we first assign scores to elements of $G$ as follows: We order the elements of $G$ according to their increasing distance to $c$ to a sequence $g_1, g_2, \ldots, g_i$. Each element of this sequence is assigned a neighborhood-normalized score

$$s_i = e^{\left(-\frac{g_i \sum i^{-1}}{\beta \sum i^{-1} g_i}\right)},$$

where $\beta$ is a parameter used to control the variance of the normalized result. From that, a set $G \times G$ of elements $g_{ij}$ is constructed with scores $s_{ij} = s_i s_j$.

At this point, each $g_{ij}$ corresponds to a pair of vertices of the SOM, which together define a 1-dimensional affine space, i.e. a line.

We use these lines to describe the position of $c$ relatively to the SOM: We orthogonally project $c$ to each line $g_{ij}$ in the multidimensional space, and obtain relative distances of the projected point $\text{proj}_{ij}(c)$ to each $g_i$ and $g_j$, as

$$d_{ij} = \frac{|\text{proj}_{ij}(c) - g_i|}{|g_j - g_i|}.$$

Distances $d_{ij}$ are used to reconstruct the embedded cell position $c'$ in the low-dimensional space space. We define the embedded SOM vertex pairs $g'_{ij} \in G' \times G'$ and relative projection distances $d'_{ij}$ accordingly for $c'$ and the chosen $G'$ (a simple choice of $G'$ may arrange the embedded SOM vertices in a lattice, but other setups are possible, see below). Finally, we aim to position $c'$ so that $d_{ij}$ and $d'_{ij}$ are as similar as possible for all $i, j$, which is accomplished by algebraically finding minimum of polynomial

$$\sum s_{ij} \|g'_i - g'_j\|^{-\alpha} (d_{ij} - d'_{ij})^2.$$

Because $d'_{ij}$ is linear in $c'$, the solution is obtained as in the least-squares method. In the formula, the neighborhood scores $s_{ij}$ provide non-linearity, and the parameter $\alpha$ is used to lower the influence of non-local information on the approximation.

As a very effective optimization, we reduce the amount of computation required for embedding all cells from the original $\mathcal{O}(|C| \cdot |G|^2)$ by truncating $G$ to $k \ll |G|$ elements that are nearest to $c$. The truncation has negligible impact on the embedding results if only elements with low $s_i$ are removed, but the total time required for embedding a set $C$ of cells is reduced to at most $\mathcal{O}(|C| \cdot |G| \cdot k^2)$.

The parameters $\alpha, \beta$ are externally represented in a more user-friendly manner. $\beta$ allows choosing the size and shape of the approximated manifold patch; we externally present it in inverse logarithmic scale as parameter Smooth that admits better interpretation by users. Similarly, parameter $\alpha$ is used to Adjust the influence of non-local information on the approximation. Effect of changing the parameters can be seen in Figure S2.

Finally, we note that the set $G'$ may be obtained in several ways. By default, EmbedSOM uses a very simple lattice layout, where $G'_{ij} = (x, y)$ if the corresponding vertex is on the $x$-th column and $y$-th row of the SOM grid. Other layouts are possible as well, examples of different layout algorithms are shown in Figure S3: 'Flat' corresponds to the default simple grid layout, 'Optimized SOM' corresponds to a layout of SOM vertices that were optimized by the Kamada-Kawai algorithm Kamada et al. [2] to better preserve distances in the U-matrix [4], 'MST based' corresponds to the vertices organized accordingly to the embedding of the minimum spanning tree of the SOM, as known from FlowSOM [5, Figure 4], the other correspond to optimization of SOM vertex positions by corresponding algorithm (UMAP and tSNE). Notably, because the size of SOM vertex set is negligible when compared to the size of whole datasets, even relatively slow embedding algorithms can be used for optimizing the layout, without any noticeable effect on performance. For example, using the Kamada-Kawai algorithm (seen e.g. in Figure 2d) for layouting added less than 1 second to the overall embedding time on all hardware we tested.

Additionally, Figure S4 shows embedding of population of developing granulocytes from Figure 1 by EmbedSOM, which is topologically correct , but overlays with other populations in the embedding and thus not easily observable.

### S3. EmbedSOM implementation and related analysis tools

The CPU-based implementation of EmbedSOM is available in the `EmbedSOM` R package, to aid interoperability with FlowSOM and other R-based software packages. Because both the SOM-building and projection stages are easily parallelizable, we also implemented GPU-accelerated versions using Vulkan® API for portability to most GPUs and platforms; the resulting package is called `vkEmbedSOM`. This accelerated implementation is based on the work of Xiao et al. [6], with modifications to make it suitable for the relatively lower dimensionality ($\leq 100$) and higher data point count ($\geq 10^6$) of cytometry data.

For the dissection, cleaning and annotation of datasets presented in the article we used `DiffSOM` R package that wraps the dissection- and statistical analysis-related FlowSOM+EmbedSOM functionality with a workflow-style interface.

All R packages are available as free software at `http://bioinfo.uochb.cas.cz/embedsom/`, along with documentation and examples of common use-cases.

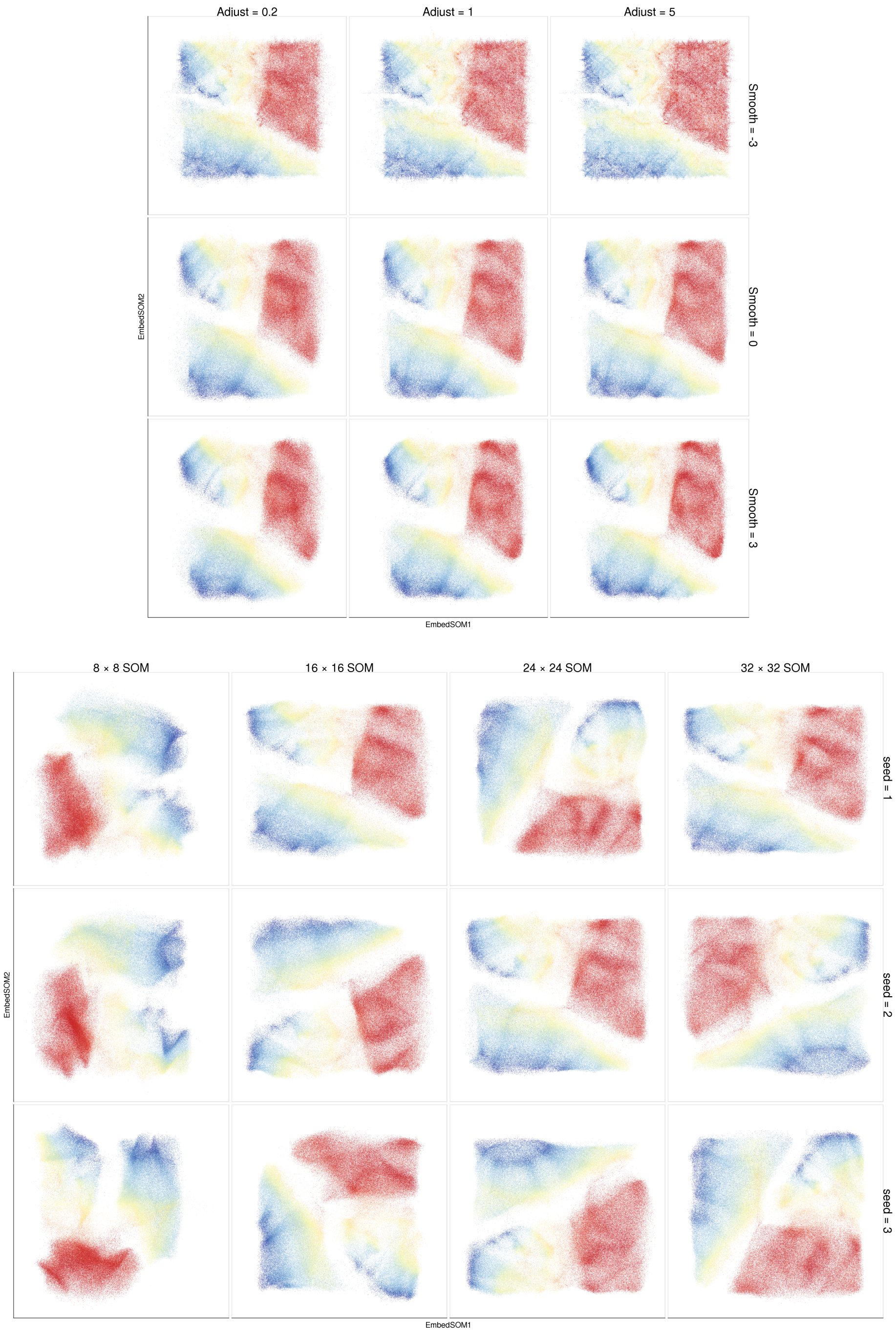


Figure S2: Effect of changing parameters demonstrated on cells from Figure 2c; cells are colorized according to CD8 expression. Top: Differences for various values of Smooth and Adjust. Bottom: Effect of changing the SOM grid size; each embedding is displayed for several different random seed values.

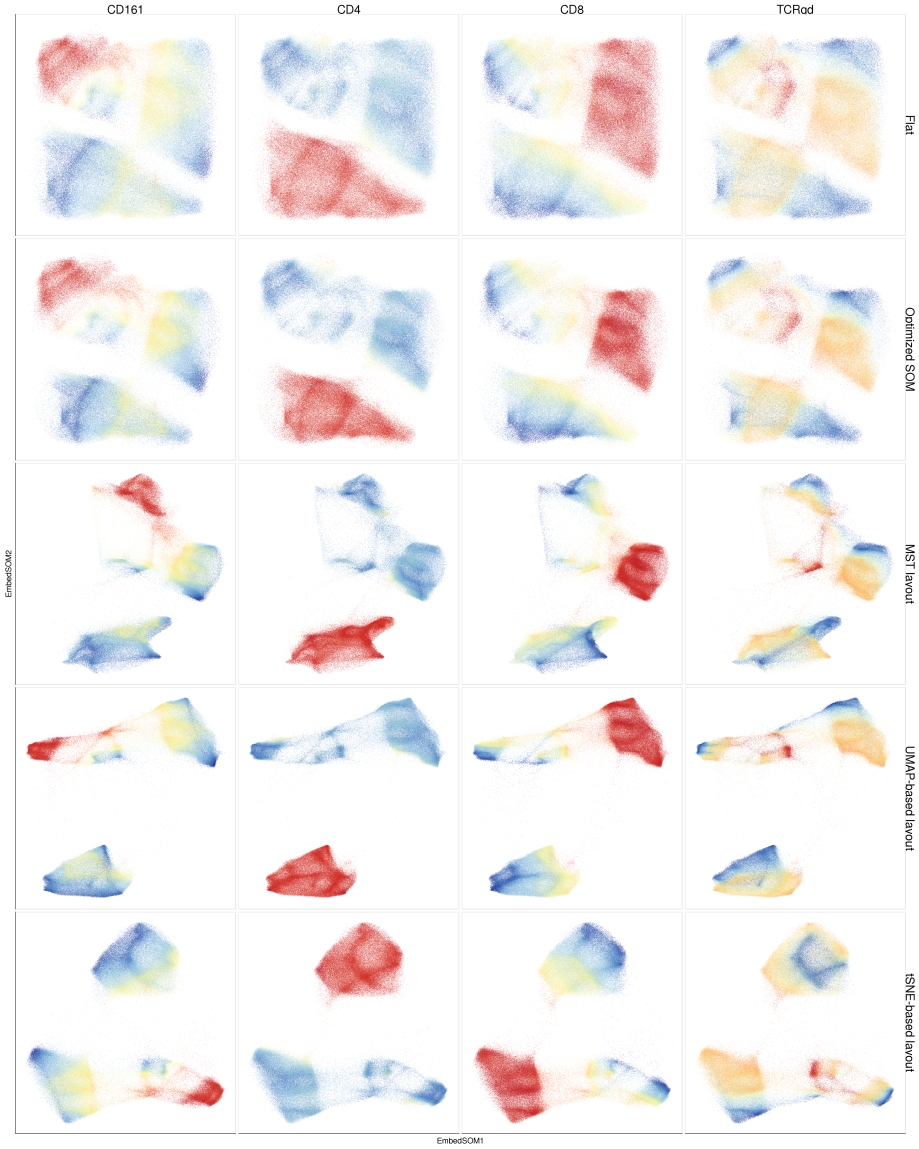


Figure S3: Comparison of various layouting possibilities for EmbedSOM; the data is the same set of cells as in Figure 2c. All layouts take approximately the same time to compute, in all cases less than 30 seconds on the testing hardware.

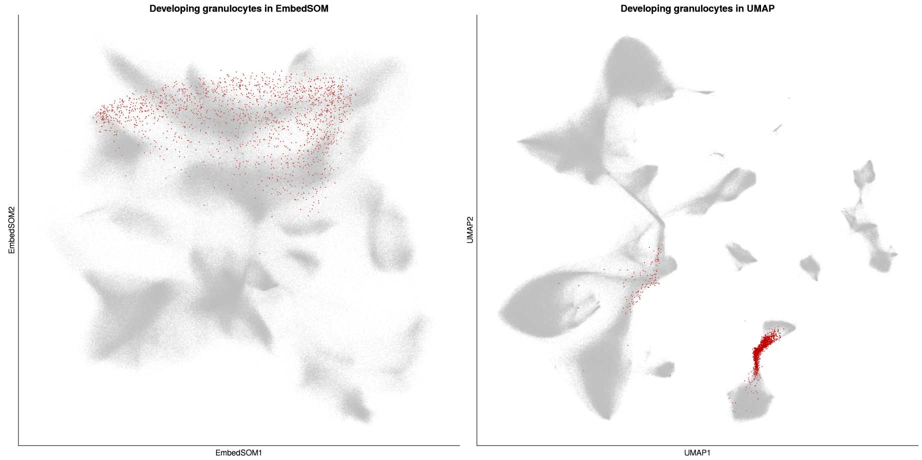


Figure S4: The cluster of developing granulocytes (highlighted in dark red) in the embedding of SamusikAll dataset in Figure 1d, using EmbedSOM (left) and UMAP (right). EmbedSOM displays it as topologically correct, as it connects the progenitor cells (upper right corner of the highlighted triangle) with eosinophils (left corner) and basophils (bottom corner) without gaps, but in overlay with other clusters and thus not clearly distinguishable. Mature neutrophils are excluded from the dataset.

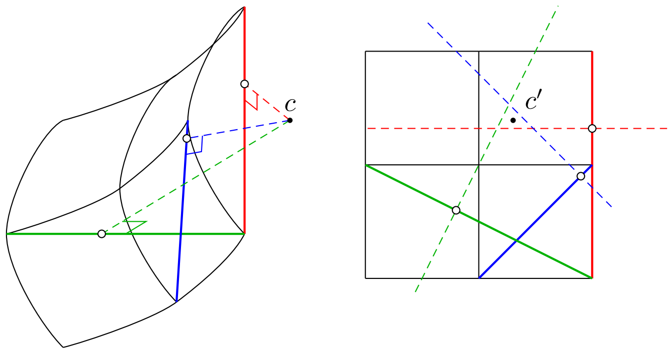


Figure S5: Approximating the projection of a single cell ($c$) in embedding stage. Left: in the multidimensional space, the cell position is described as a collection of its projections into 1-dimensional affine spaces defined by pairs of SOM clusters. Right: the same situation is trivially reconstructed in 2 dimensions; the final cell position is fitted to minimize the error in the projection coordinates.

| dataset | #events | used dims. | used cells | annotated populations | FlowRepository |
|---|---|---|---|---|---|
| Levine13 | 167k | 13 | 82k | 24 | FR-FCM-ZZPH [3] |
| Levine32 | 265k | 32 | 104k | 14 | |
| Samusik01 | 86k | 39 | 53k | 24 | |
| SamusikAll | 841k | 39 | 514k | 24 | |
| Pregnancy | 90.2M | 41 | 23.9M | – | FR-FCM-ZY3Q [1] |
| Mouse dissection | 4.8M | 14 | 1M | – | available online[2] |

Table S1: Summary of the publicly available datasets used for demonstrations and benchmarking.

### S4. Benchmarks and benchmarking datasets

The data used for benchmarking were obtained from publicly available datasets; a summary is available in Table S1. The embeddings used in benchmarking are displayed in Figure S6, together with embeddings with other EmbedSOM layout methods.

The whole benchmark methodology was set up to be easily replicated using standardized tools and processing; code required for generating all results in the paper is available on GitHub at `https://github.com/exaexa/EmbedSOMArticle/` (currently available upon request).

The repository is organized as a set of scripts that are run in workflow hierarchy according to `Makefile`-defined recipes. After invoking a complete build, this description triggers download of the original datasets, recomputation of all benchmark results and replication of all analyses from raw data, regeneration of all data-derived figures used in the article, and typesetting a complete LaTeX version of this article. We hope that such presentation of the workflow will improve the reproducibility of the results and re-usability of the methods involved.

[2] Mouse dissection data are available from `https://bioinfo.uochb.cas.cz/embedsom/datasets/C067_f.tar.xz`.

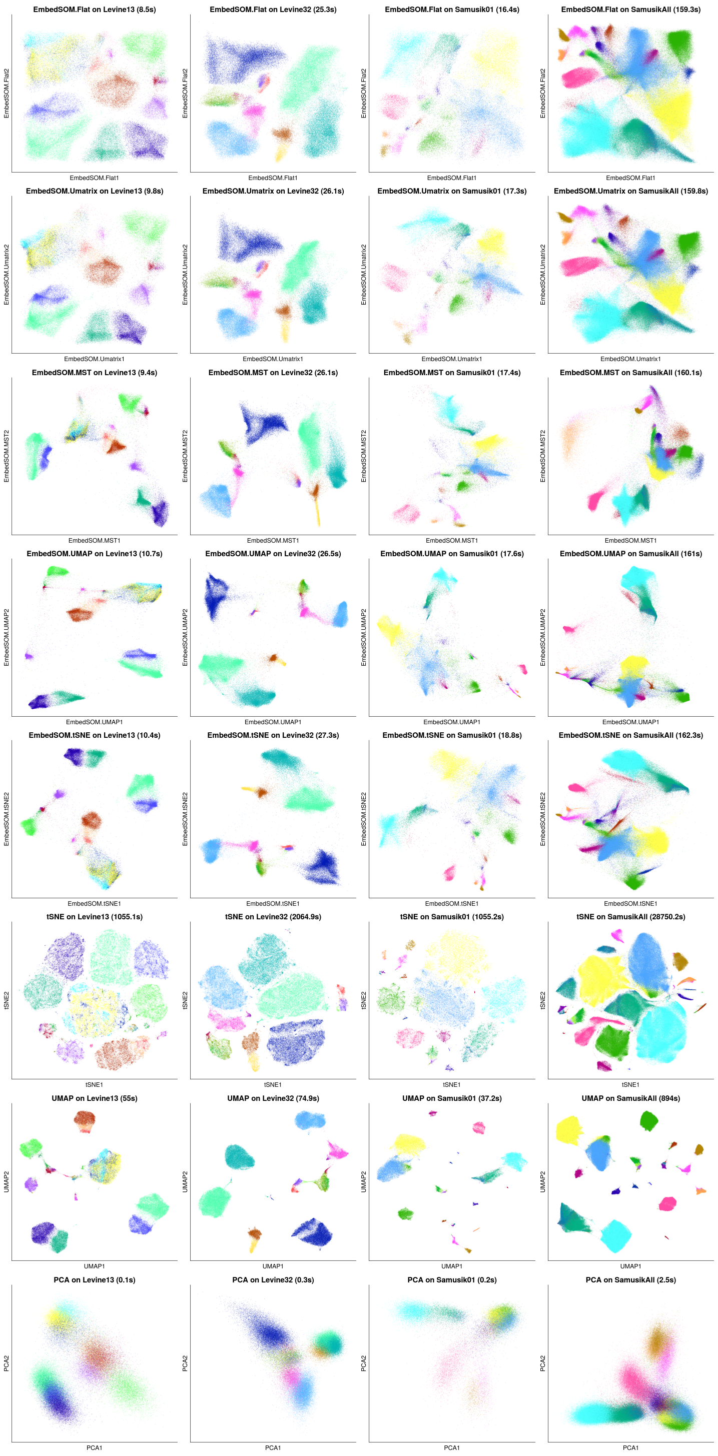


Figure S6: Various embeddings of the Levine13, Levine32, Samusik01 and SamusikAll datasets. The datasets were prepared as described in Section 4.1.

### References


[1] Aghaeepour, N., Ganio, E.A., Mcilwain, D., Tsai, A.S., Tingle, M., Van Gassen, S., Gaudilliere, D.K., Baca, Q., McNeil, L., Okada, R., et al., 2017. An immune clock of human pregnancy. Science immunology 2, eaan2946.

[2] Kamada, T., Kawai, S., et al., 1989. An algorithm for drawing general undirected graphs. Information processing letters 31, 7–15.

[3] Levine, J.H., Simonds, E.F., Bendall, S.C., Davis, K.L., ad D. Amir, E., Tadmor, M.D., Litvin, O., Fienberg, H.G., Jager, A., Zunder, E.R., Finck, R., Gedman, A.L., Radtke, I., Downing, J.R., Pe'er, D., Nolan, G.P., 2015. Data-driven phenotypic dissection of AML reveals progenitor-like cells that correlate with prognosis. Cell 162, 184–197.

[4] Ultsch, A., 2003. U*-matrix: a tool to visualize clusters in high dimensional data. Technical report, published by Fachbereich Mathematik und Informatik Marburg.

[5] Van Gassen, S., Callebaut, B., Van Helden, M.J., Lambrecht, B.N., Demeester, P., Dhaene, T., Saeys, Y., 2015. FlowSOM: Using self-organizing maps for visualization and interpretation of cytometry data. Cytometry Part A 87, 636–645.

[6] Xiao, Y., Feng, R.B., Han, Z.F., Leung, C.S., 2015. GPU accelerated self-organizing map for high dimensional data. Neural Processing Letters 41, 341–355.
